# Towards a taxonomically unbiased EU Biodiversity Strategy for 2030

**DOI:** 10.1101/2020.07.06.189027

**Authors:** Stefano Mammola, Nicoletta Riccardi, Vincent Prié, Ricardo Correia, Pedro Cardoso, Manuel Lopes-Lima, Ronaldo Sousa

**Author notes:** **Author contribution:** MLL, NR, RS, SM, and VP conceived the idea. NR mined data from LIFE projects. RC extracted data from IUCN and the cultural value of species. NR, RS, and SM extracted species body size from different sources. SM performed analyses and prepared figures. RS and SM wrote the first draft of the paper. All authors critically contributed to the writing of the paper through comment, additions, and revisions.

## Abstract

Through the Habitats Directive (92/43/EEC) and the LIFE projects financial investments, Europe has been the world’s experimental arena for biological conservation. With an estimated budget of €20 billion/year, the EU Biodiversity Strategy for 2030 has set an ambitious goal of reaching 30% Protected Areas and ensure no deterioration in conservation trends and status of all protected species. We analyzed LIFE projects focused on animals from 1992 to 2018 and we found that investment towards vertebrates has been six times higher than that for invertebrates (€970 vs €150 million), with birds and mammals alone accounting for 72% species and 75% total budget. Budget allocation is primarily explained by species’ popularity. We propose a roadmap to achieve unbiased conservation targets for 2030 and beyond.

## Main Text

Overwhelming evidence exists that most Earth ecosystem processes are now altered by human activities, suggesting that we may have entered a human-dominated geological epoch – the ‘Anthropocene’ (Lewis and Maslin 2015). It is largely accepted that humans are causing the sixth mass species extinction (Dirzo et al. 2014), which can be considered a clarion call to increase global efforts to study, halt, and possibly reverse the ongoing negative environmental trends. Europe is no exception given that it has a long experience of human disturbance and consequent biodiversity loss (Müller et al. 2020). At the same time, since the Habitats Directive (92/43/EEC) was established in 1992, the European Union (EU) has also been the world’s experimental arena for practical conservation and restoration of natural habitats and their wild flora and fauna.

Although the Habitats Directive and the parallel financial investment on LIFE projects – i.e., the EU flagship funding instrument for the environment and climate action created in 1992 – are seen as pioneer endeavors and represent a strong legal and financial tool for biodiversity conservation, the number of species in Europe continues to decline (Müller et al. 2020; but see Chapron et al. 2014 for positive rewilding trends in a few charismatic large carnivores). At the end of the trial period of the Habitats Directive, the EU has launched a new Biodiversity Strategy intending to achieve 30% of Protected Areas by 2030 and ensure no deterioration in conservation trends and status of all protected species and habitats (European Commission 2020). The ideals of this ambitious plan resonate with that of other similar projects such as the Global Deal for Nature (Dinerstein et al. 2019) or the Half-Earth project (E.O. Wilson Biodiversity Foundation), aiming to protect up to 50% of the Earth’s ecoregions. Such an effort will be critical if we are to embrace the recent proposal of keeping described species extinctions to below 20 a year over the next 100 years (Rounsevell et al. 2020).

These conservation schemes should ideally encompass all native ecosystem types, species, and ecological successions, in order to ensure not to *“[…] add more land to reach the global target that is similar to what is already well accounted for at the expense of underrepresented habitats and species”* (Dinerstein et al. 2019). It is therefore mandatory to assess whether conservation investment is optimally allocated among species and habitats, or taxonomic and other biases still permeate biological conservation (Clark and May 2002). As the EU is proposing a large budget to be annually unlocked for spending on nature (estimated €20 billion/year; European Commission 2020), the time is ripe to take stock of what the Habitats Directive and LIFE projects have achieved and look forward to the goal of establishing an unbiased conservation agenda in Europe.

### Allocation of projects and fundings

To quantify the past taxonomic biases and potential drivers over a long-term scale during which they can potentially impact conservation strategies and decision-making policies, we mined information on LIFE projects focused on animals carried out between 1992 and 2018 (n = 835). With this exercise, our goal was to obtain a comprehensive picture about the number of applied conservation actions and allocation of the EU’s conservation budget across the animal Tree of Life in Europe (full methodological details are in Supplementary material Appendix S1).

The number of projects (Fig. 1a) and budget allocation (Fig. 1b) have varied substantially across faunal groups. Overall, LIFE projects focused on 410 vertebrate and 78 invertebrate species. Net monetary investment towards vertebrates was over six times higher than that for invertebrates (*ca.* €970 vs €150 million; Table S1), a long-known taxonomic bias (Clark and May 2002; Fukushima et al. 2020) that seemingly keeps operating in conservation throughout Europe (Mammides 2019). This disproportion in conservation efforts is even more striking when relativizing it to the actual numbers of known species in Europe, namely *ca.* 1,800 ssp. of vertebrates and 130,300 spp. of invertebrates (de Jong et al. 2014). In relative terms, 23% of vertebrates received funding versus only 0.06% of invertebrates and the total investment per species towards vertebrates has been 468 times higher than that for invertebrates. Not counting the species of invertebrates as yet to be discovered (Stork 2018), including in Europe.

**Fig. 1:**
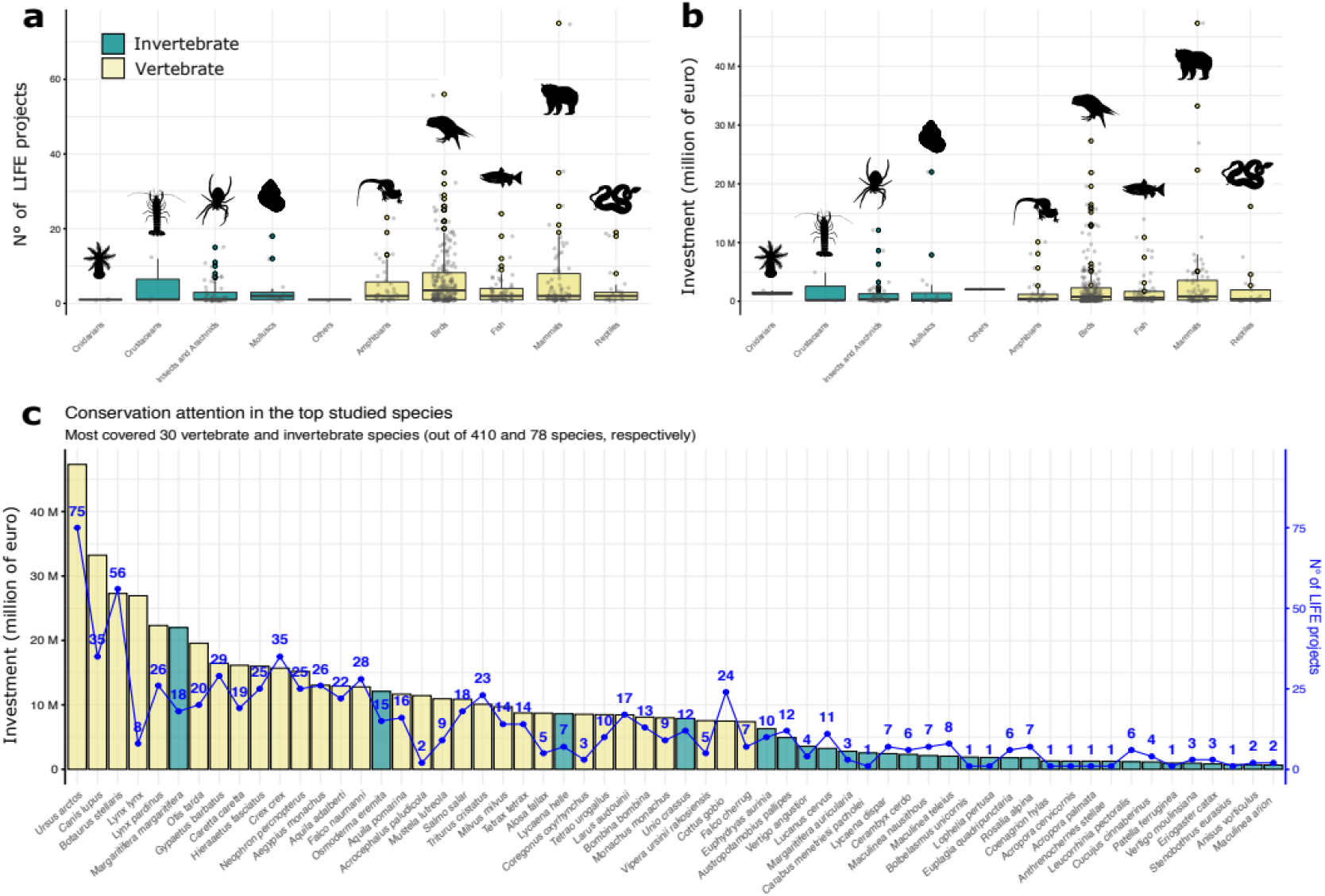
**a–b)** Breakdown of number of projects **(a)** and budget allocation **(b)** across main animal groups covered by the LIFE projects (n = 835). **c)** Most covered 30 species of vertebrates (out of 410) and invertebrates (out of 78) by the LIFE projects analysed (n = 835). Vertical bar is monetary investment and blue scatter line number of LIFE projects devoted to each species (see Fig. S1 for the graphs of species alone).

The top 30 invertebrate species, regarding LIFE funds allocation (Fig. 1c), are coleopterans, butterflies, dragonflies, and a few molluscs; a drop in the ocean given invertebrate diversity and current rate of declines across diverse habitats (Eisenhauer et al. 2019; Cardoso et al., 2020). Even within invertebrates the biases are notorious with widespread, large, and/or colourful species being dominant (Cardoso 2012). Among vertebrates, 54% of species covered by LIFE projects were birds (accounting for 46% of the budget allocated) and 18% mammals (24% of the budget), with only 8 out of 30 most protected vertebrate species not belonging to these two groups (Fig. 1c).

As expected, we found a significant positive correlation between the allocation of budget and the number of projects (Fig. 2a), although with some differences in conservation strategies for distinct species. Whereas the majority of species were covered by few projects with a limited budget (in the range €0– 5 million), some outlying species were the subject of more intense budget allocation and several projects (e.g., bear and great bittern), or of a lower number of high-budget projects (e.g., lynx and wolf). The only outlier among invertebrates was the freshwater pearl mussel *Margaritifera margaritifera,* being the focus of a few high-budget projects (Fig. 1C and 2a). It should be noted, however, that all these species are distributed over broad geographic expanses encompassing several EU countries, and thus are more prone to be targeted by researchers applying to LIFE projects. Ironically, these broad geographic ranges possibly turn these species less prone to extinction.

**Fig. 2:**
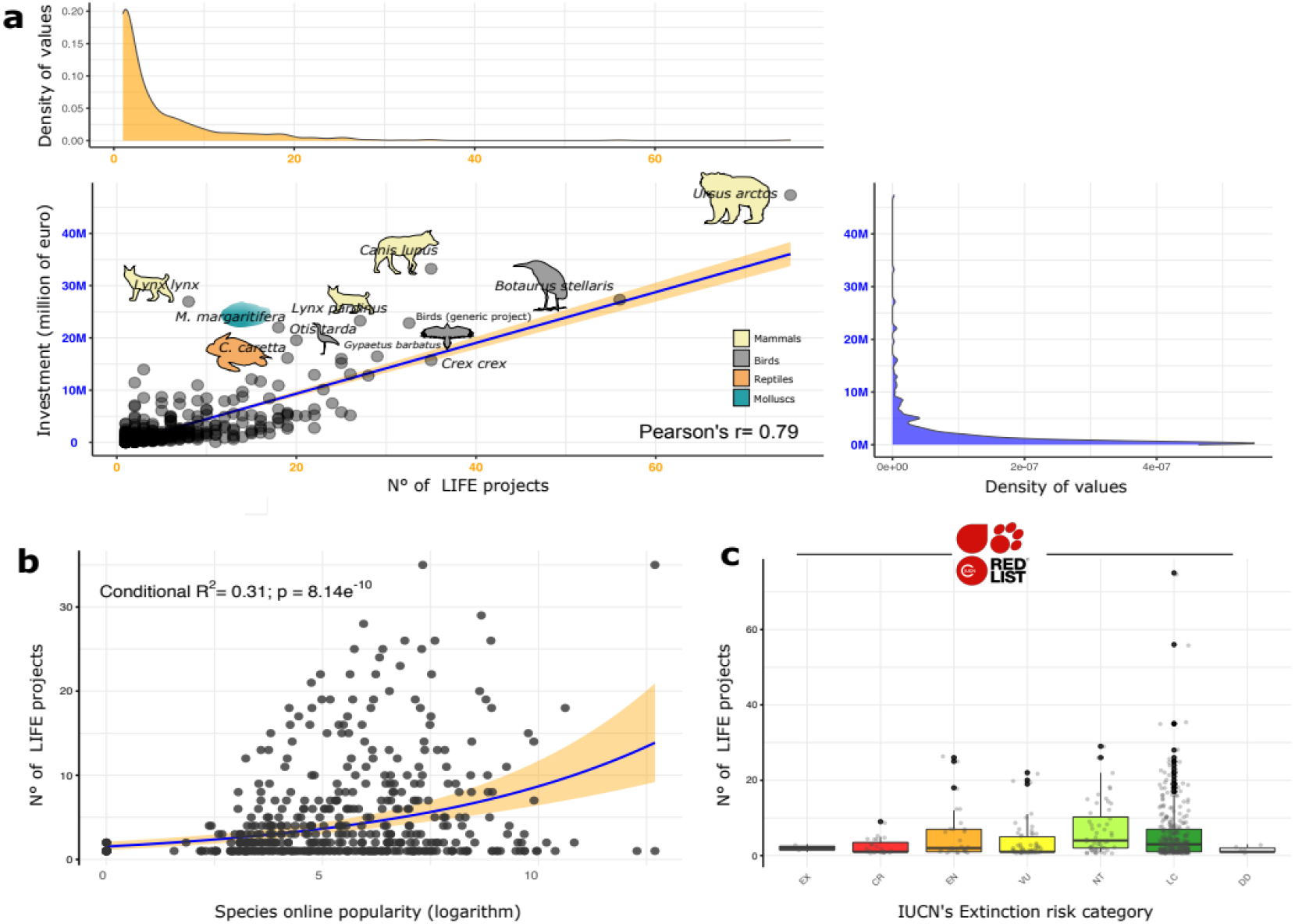
**a**) Pearson’s *r* correlation between monetary investment and number of projects for the 78 invertebrate and 410 vertebrate species covered by LIFE projects (n = 835). The farther away a dot is from the correlation line, the more the conservation effort is unbalanced toward either the number of projects or the monetary investment. **b**) Predicted relationship (blue line) and 95% confidence interval (orange shaded surface) between the number of LIFE projects and the species’ online popularity, according to the results of a negative binomial Generalized linear mixed model (full details on the analysis in Appendix S1). Online popularity is measured as the net online attention each species receives. **c)** Breakdown of LIFE projects according to the IUCN extinction risk of the species that they cover (EX: Extinct; CR: Critically Endangered; EN: Endangered; VU: Vulnerable: NT: Near Threatened; LC: Least Concern; DD: Data Deficient).

### Drivers of taxonomic bias

To obtain a deeper understanding of the factors underlying the allocation of conservation investment among species which were accounted for by LIFE projects, we asked whether the taxonomic bias has been due to:

i. an objectively higher extinction risk – approximated using International Union for Conservation of Nature’s (IUCN) extinction risk categories;
ii. the conspicuousness of species – approximated as species body size, one of the few comparable traits among so diverse taxa and also a putative driver of extinction risk (Chichorro et al. 2019, 2020, Carmona et al. 2020); or
iii. cultural reasons – approximated as the online popularity of each species which we measured using a culturomics approach (Ladle et al. 2016).

We expressed conservation investment simply as the number of LIFE projects, given that the number of projects was highly correlated with total budget (Fig. 2a). We explored the contribution of IUCN extinction risk, body size, and people’s interest in driving conservation investment in species that received funding with a generalized linear mixed model (GLMM) that took into account taxonomic relatedness among species (Appendix S1). We found that the only significant predictor explaining the conservation attention a species receives is online popularity (negative binomial GLMM: estimated β ± S.E.: 0.18 ± 0.03; p < 0.001; Table S2). In particular, species being covered by a greater number of LIFE projects were also those attracting most interest online (Fig. 2b), suggesting that conservation in the EU is largely driven by species charisma, rather than objective features. This result aligns with a recent study recovering a similar trend in a global sample of threatened vertebrates (Davies et al. 2018).

Unexpectedly, the species risk of extinction does not seem to affect LIFE funds allocation given that species under extinction threat categories did not receive significantly higher conservation attention. On the contrary, we found that the greater number of projects focused on Least Concern and Near Threatened species (Fig. 2c). Over the 26 years of LIFE projects there have been strong disagreements between the Red Lists and the Habitats Directive protected species list, which would call for careful revision of the EU policy (for further discussion see Moser et al. 2016). Whereas some of the Habitats Directive species that are not included in threatened IUCN categories may be broadly distributed thereby acting as umbrella species for the habitat that they occupy and the species they coexist with, this disproportion of investment seems unjustified given the limited budget of past LIFE projects.

The non-significant effect of body size was also somewhat unexpected given that this trait is often a good proxy of species’ extinction risk, although it depends on the taxon (Chichorro et al. 2020). However, it should be noted that most invertebrates in the Habitats Directive are large species. Moreover, the correlation between people’s interest and body size was quite high (Fig. S4) and thus, the strong and significant effect of people’s interest may have partly masked a weaker effect of body size in explaining conservation attention.

### A way forward

The commitment of the EU to realize the ambitious goals of the Biodiversity Strategy for 2030 appears evident when considering the proposed financial plan for nature conservation (European Commission 2020). At least €20 billion a year will be unlocked for spending on nature, which will require mobilising private and public funding at national and EU level, including through a range of different programs. Moreover, as nature restoration will make a major contribution to climate change mitigation objectives, a significant proportion of the 25% of the EU budget dedicated to climate action will be invested on biodiversity and nature-based solutions. All these actions go hand-in-hand with the EU Green deal that emphasizes the post-covid-19 economic recovery, with the intention to further invest in conservation and sustainable development. However, if we look at past investments taking the Habitats Directive and LIFE projects as a proxy, we may predict that a few charismatic species will receive almost all conservation attention. In targeting 30% of protected areas and to halt the documented trends of species extinctions (Rounsevell et al. 2020), especially the silent extinctions of invertebrates (Eisenhauer et al. 2019), it is mandatory to overcome prevalent taxonomic biases. We hereby propose a roadmap for the next decade.

First, we need to invest in species inventory and data compilation, to overcome main knowledge gaps (discussed in Cardoso et al. 2011). Aside from funding, overcoming gaps will require creativity in making use of diverse sources of data, including monitoring schemes that already exist at the national (e.g., U.K. and Germany) and EU levels (e.g., Water Framework Directive monitoring for freshwater taxa), citizen science projects (e.g., BioBlitz), and internet-derived data (*iEcology*; Jarić et al. 2020).

Second, armed with such knowledge, we should quickly assess the status of a broader sample of species to obtain a real picture about the fraction of diversity that is threatened by human activities. Currently only a handful of invertebrates are included in the IUCN red list, preventing us from fully understanding their conservation status and trends (Cardoso 2012).

Third, it is critical to review the Habitats Directive Annexes considering unbiased criteria for species protection. This will allow us to build sound scenarios of possible enlargement of the Natura 2000 network to implement 30% coverage in the EU member states’ protected areas (see Müller et al. 2020).

Fourth, we should focus on endangered species and their habitats rather than directing substantial conservation efforts towards non-threatened species (Fig. 2c). The EU Biodiversity Strategy for 2030 will request the Member States to ensure no deterioration in conservation trends and status of all protected habitats and species by 2030. This goal will be reached only by directing conservation efforts to species at greater risk of extinction, and by focusing on those taxa and habitats that are currently not accounted for (Dinerstein et al. 2019).

Fifth, going forward, it will be necessary to monitor conservation priority and funding investment on an annual basis. In the EU there is no comprehensive governance framework to steer the implementation of biodiversity commitments agreed at the national, European, or international level. To address this gap, the Commission is planning a new European biodiversity governance framework to map obligations and commitments and set out a roadmap to guide their implementation. As part of this new framework, a monitoring and review mechanism will be established, including a clear set of agreed indicators. It is within this framework that we must challenge the current taxonomic bias and establish a more equitable redistribution of funds.

Considering the number of European species we are dealing with (de Jong et al. 2014), we realize that it will not be possible to implement this agenda for all species and habitats. As proposed by Cardoso & Leather (2019), the simplest solution will be to maximize phylogenetic and functional coverage of the species targeted for the LIFE projects. An easy-to-implement approach would be to include a positive discrimination mechanism by which sound LIFE projects focusing on neglected threatened species will be given priority. This can be achieved by weighting future project review scores according to objective criteria based on phylogenetic, functional, and spatial uniqueness in relation to previous projects. This way, species that have received substantial funding in the past (e.g., bear and lynx) would see their scores down-weighted; conversely, species that have never received funds (e.g., most invertebrates) would increase their project score. These, or similar approaches and criteria, will increase objectivity of future EU conservation planning, leading the world by example and by action.

## Supplementary Materials

**Appendix S1**: Extended Methods.

**Table S1:** LIFE projects investment breakdown among animal groups.

**Table S2:** Estimated regression parameters for the negative binomial Generalized linear mixed model.

**Fig. S1:** Most covered invertebrates and vertebrates species by the LIFE projects analyzed.

**Fig. S2:** Multicollinearity among continuous variables.

**Fig. S3:** Association between continuous and categorical variables.

**Fig. S4:** Relationship between body size and species’ public interest.

## Data availability statement

Data supporting this study and R code to generate graphs and analyses will be made freely available in Figshare upon acceptance in an official peer-reviewed journal.

## Supplementary material

### Supplementary items

**Appendix S1**: Extended Methods.

**Table S1**: LIFE projects investment breakdown among animal groups.

**Table S2**: Estimated regression parameters for the negative binomial Generalized linear mixed model.

**Figure S1**: Most covered invertebrates and vertebrates species by the LIFE projects analyzed.

**Figure S2**: Multicollinearity among continuous variables.

**Figure S3**: Association between continuous and categorical variables.

**Figure S4**: Relationship between body size and species’ public interest.

## S1. EXTENDED METHODS

### Extraction of data from LIFE project

We extracted information on the amount of funding allocated to the various species using the LIFE projects database (https://ec.europa.eu; accessed between February and May 2020). We first filtered LIFE projects specifically aimed at species conservation, using the query STRAND = “All”; YEAR = “All”; COUNTRY = “All”; THEMES = “Species”; SUB-THEMES = “Amphibians”; “Birds”; “Fish”; “Invertebrates”; “Mammals”; “Reptiles”. This query resulted in 819 LIFE projects that meet search criteria. A second query focused on THEMES = “Biodiversity issues”; SUB-THEMES = “Ecological coherence”; “Invasive species”; “Urban biodiversity”; matched an additional 298 LIFE projects. For the latter query, we examined summaries of the LIFE projects and extracted further information only from those specifically aimed at the conservation of animal species (n = 16). For example, projects aimed at generically enhancing biodiversity through actions on ecosystems or impacts caused by anthropic activities were not considered. In total, we included 835 projects in the analyses – 819 with theme “Species” and 16 with theme “BIodiversity issues”. To define the amount of funds allocated to each species for each LIFE project, the budget of each project with multiple species was redistributed equally among the target species.

### Calculation of species traits

To obtain a deeper understanding of the factors underlying the observed pattern of allocation of conservation effort among species, we asked whether the number of LIFE projects and the budget allocation for each species is driven by its risk of extinction (IUCN 2020), body size, and/or the online popularity of the species (Davies et al. 2018). We measured these features using three proxies, as detailed in the following sections.

#### Species extinction risk

We estimated the extinction risk category of each species in our database using the International Union for Conservation of Nature Red List of Threatened Species (IUCN 2020). Using API keys, we automatically assigned each species to its IUCN extinction risk category and then, we manually checked this matching to correct potential mistakes.

#### Body size

Body size is one of the few traits that is fully comparable across distant taxa (Weiss and Ray 2019) and also, it is one of the most conspicuous traits correlating with extinction risk (see also Fig. S3a). We estimated body size for birds using the Collins birds guide (Svensson et al. 2010), for mammals using Aulagnier et al. (2018), for reptiles and amphibians with Speybroeck et al. (2016), for fish using Fishbase (Froese and Pauly 2000). For invertebrates, we estimated body size using diverse sources, from Wikipedia to original species descriptions and expert opinion and unpublished data by the authors (e.g. molluscs or insects).

#### Cultural salience of each species

We characterised the online popularity of each species in our database using a culturomics approach (Ladle et al. 2016) based on the volume of internet searches carried out through Google’s Search Engine. We obtained data on the average monthly relative search volume recorded between January 2010 and December 2019 for each species from the Google Trends API. Google Trends returns relative search volume data ranging from 0 to 100; the maximum value is assigned to the highest proportion of total searches observed during any given month of the sampled period and all other values are scaled concerning it. To ensure comparable data between species, we collected the data using an approach similar to that described by Davies et al. (2018). We carried out topic searches for multiple species combinations, validating each species topic beforehand (for details see Correia 2019), and ensuring one common species between different searches. The values returned for this species in either search were used to estimate a scaling factor between searches, calculated as the coefficient of a linear regression between the values of both searches. The species used to calculate the scaling factor was selected iteratively to ensure the scaling factor was calculated as accurately as possible based on i) the highest number of non-zero values between searches, and ii) a regression R2 value above 0.95.

### Statistical analyses

To obtain insight on which specific traits (IUCN extinction risk, body size, and/or people interest) are driving conservation attention, we explored the relationships between species traits and LIFE project funding with generalized linear mixed model that accounted for taxonomic non-independence among species (Zuur and Ieno 2016). Given that the number of projects and budget allocation were highly correlated (Pearson’s *r* = 0.8; Fig. S2), we only selected the number of projects as a proxy measure for conservation attention.

We performed analyses in R (R Core Team 2018). We initially explored the dataset following a general protocol for data exploration (Zuur et al. 2009a). We checked homogeneity of continuous variables and log-transformed non-homogeneous variables, when appropriate. We verified multicollinearity among predictors with pairwise Pearson’s *r* correlations (Fig. S2), setting the threshold for collinearity at |*r*| > 0.7 (Zuur et al. 2009b). We visualized potential associations between continuous and categorical variables with boxplots (Fig. S3). Given that average people interest (logarithm) and relative people interest (logarithm) were highly correlated (*r* = 0.96), we only included average people interest in the regression analysis.

We fitted a Generalized linear mixed model (GLMM) to the data with the R package *lme4* (Pinheiro et al. 2019). The mixed part of the model allowed us to take into account taxonomic non-independence of observations, under the assumption that closely related species should share more similar traits than expected from random. Specifically, the taxonomic relatedness among species was accounted for with a nested random intercept structure (Class / Order / Family). Given that the response variable number of projects are counts, we initially selected a Poisson error and a log link function. The Poisson model was, however, slightly over-dispersed (dispersion ratio = 3.5; Pearson’s χ^2^ = 1463.5; *p*-value < 0.001) and thus, we switched to a negative binomial distribution. The estimated regression parameters are in Table S1. Model validation was performed with the aid of the R package *performance* (Lüdecke et al. 2020).

## Supplementary Tables

**Table S1:** LIFE projects investment breakdown among animal groups (1992–2018; n = 835).

| Animal group | LIFE project funding allocation [Euro] |
| --- | --- |
| Amphibian | 53,424,690.98 |
| Birds | 447,771,421.63 |
| Fish | 135,258,976.43 |
| Mammals | 387,716,614.01 |
| Reptiles | 46,103,756.32 |
| Invertebrates | 154,577,014.64 |

**Table S2:** Estimated regression parameters for the negative binomial Generalized linear mixed model. S.E. = standard error; EX: Extinct; CR: Critically Endangered; EN: Endangered; VU: Vulnerable: NT: Near Threatened; LC: Least Concern; DD: Data Deficient. Signif. codes = (*): Approaching significance; *: Significant.

| Variable | Estimated $\beta \pm \text{S.E.}$ | *p*-value |
| --- | --- | --- |
| Intercept (IUCN baseline: CR) | 0.53 ± 0.33 | - |
| IUCN – EX | 0.05 ± 1.03 | 0.95 |
| IUCN – EN | 0.36 ± 0.33 | 0.27 |
| IUCN – VU | -0.03 ± 0.31 | 0.90 |
| IUCN – NT | 0.53 ± 0.31 | 0.07 (*) |
| IUCN – LC | 0.17 ± 0.28 | 0.55 |
| People interest (logarithm) | 0.18 ± 0.03 | < 0.001 * |
| Body size (logarithm) | -0.07 ± 0.05 | 0.17 |

## Supplementary Figures

**Fig. S1:**
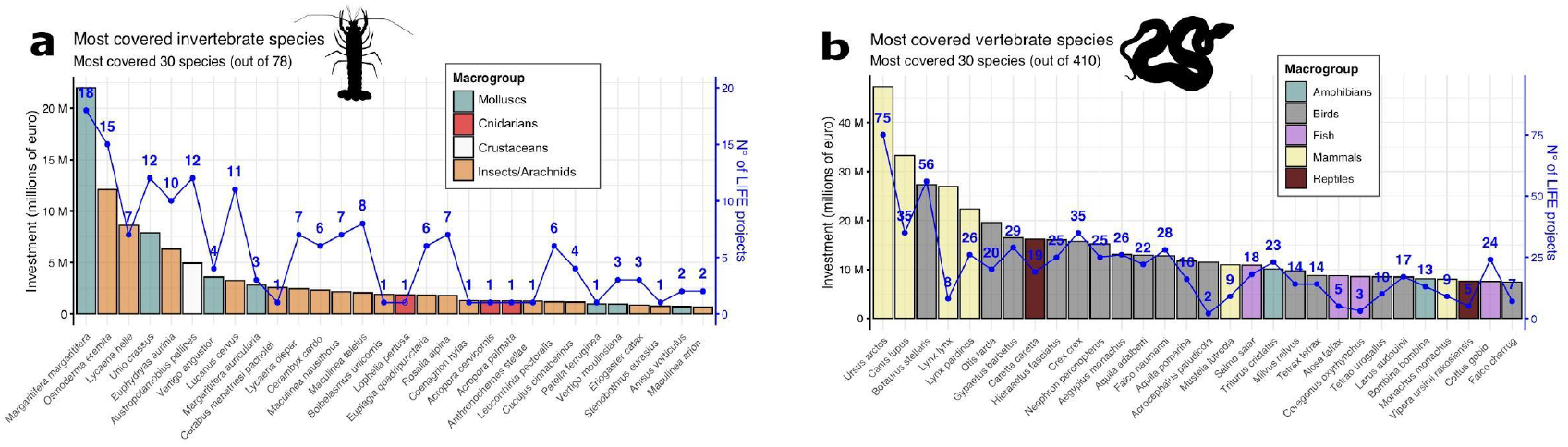
Most covered 30 species of (**a**) invertebrates (out of 78) and (**b**) vertebrates species by the LIFE projects analyzed (n = 835). Vertical bar is monetary investment and blue scatter line number of LIFE projects devoted to each species.

**Fig. S2:**
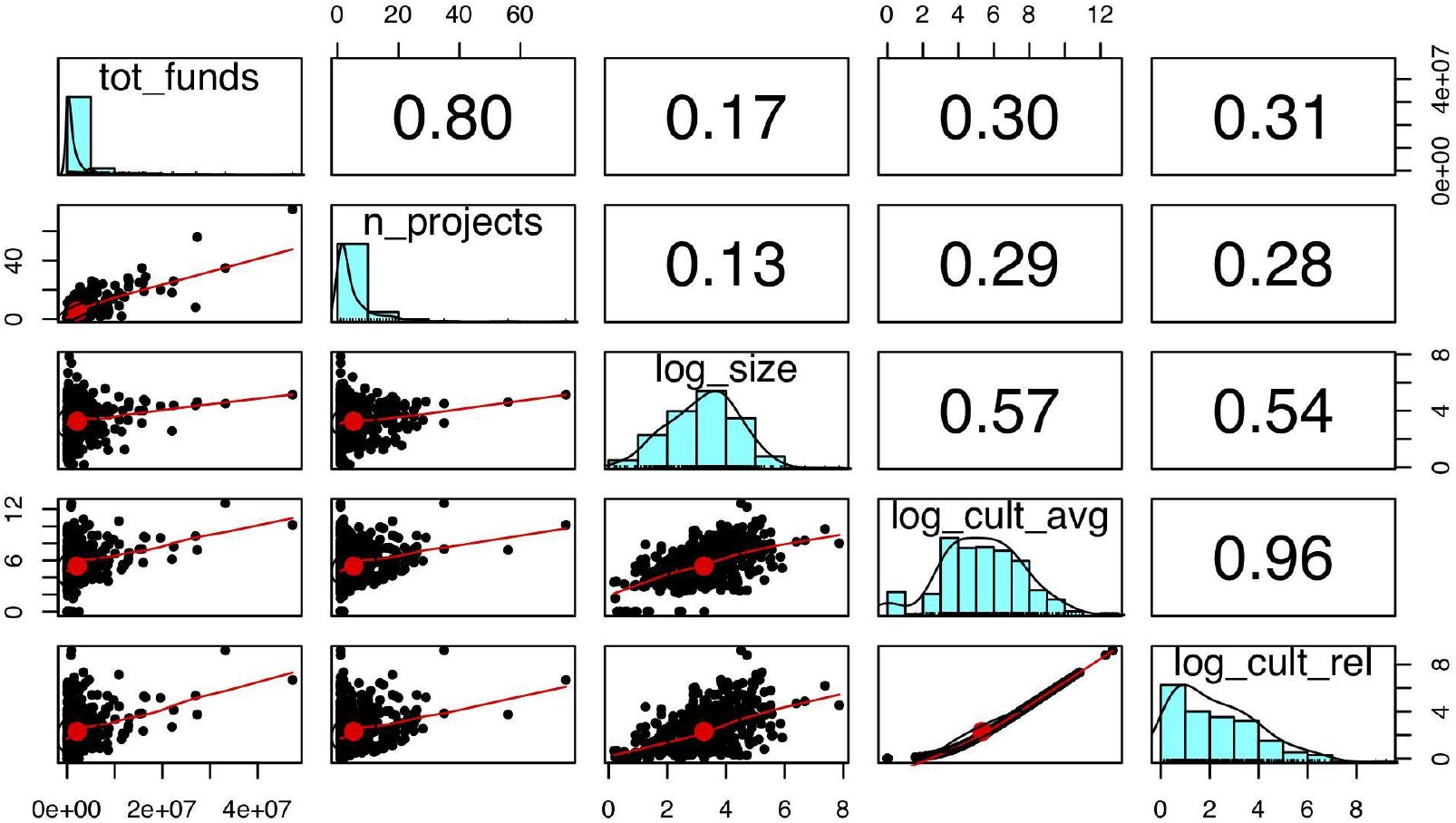
Multicollinearity among continuous variables. Above the diagonal: Pearson *r* correlation coefficient. On the diagonal: histogram illustrating data distribution. Above the diagonal: scatter plot of x by y with a trendline fitted through the data. tot_funds: total monetary investment; n_projects: N° of LIFE projects; log_size: body size (logarithm); log_cult_avg: average people interest (logarithm); log_cult_rel: relative people interest (logarithm).

**Fig. S3:**
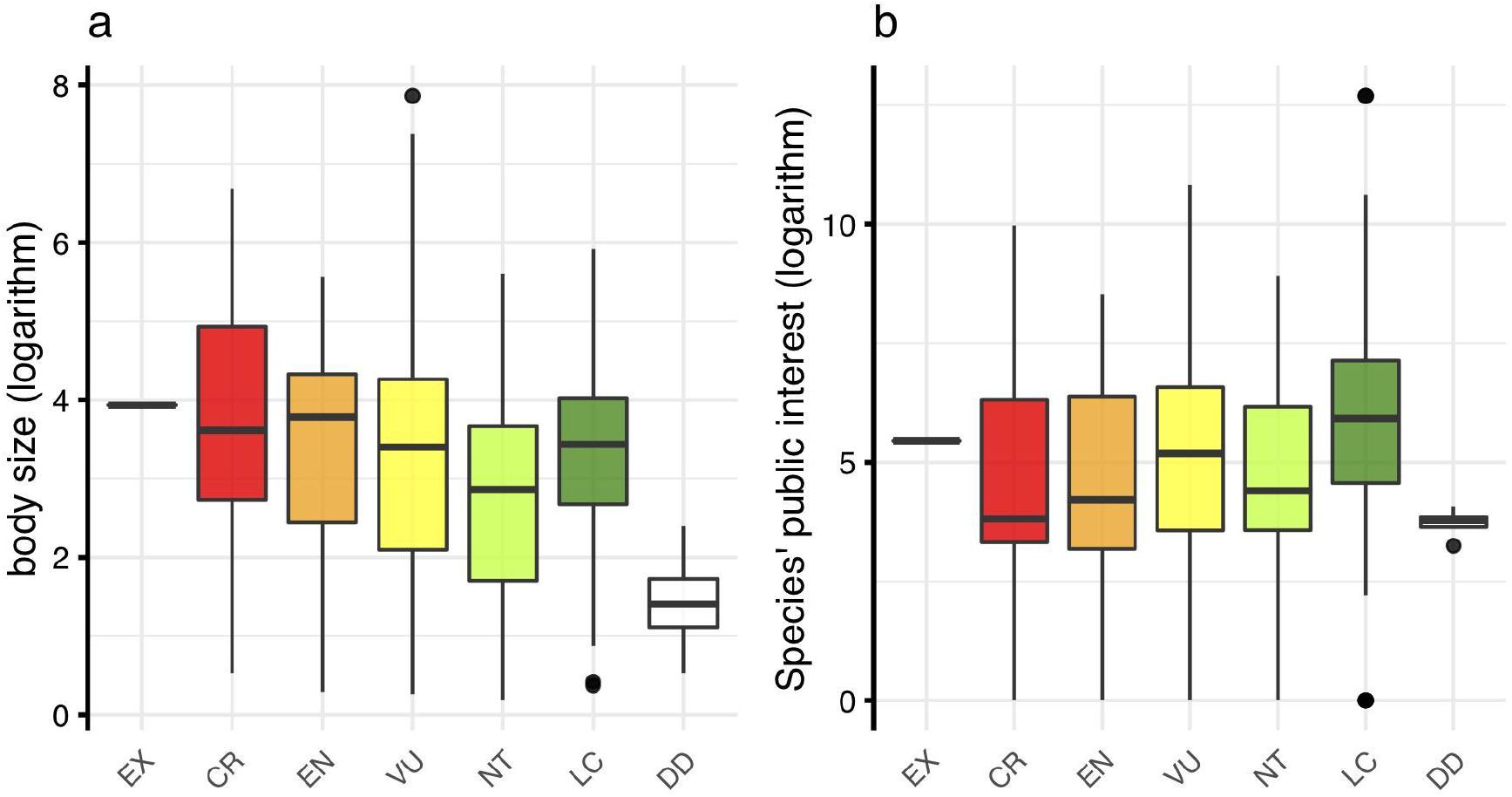
Association between continuous and categorical variables used in the regression models. **a**) Body size **b**) Species’ online popularity.

**Fig. S4:**
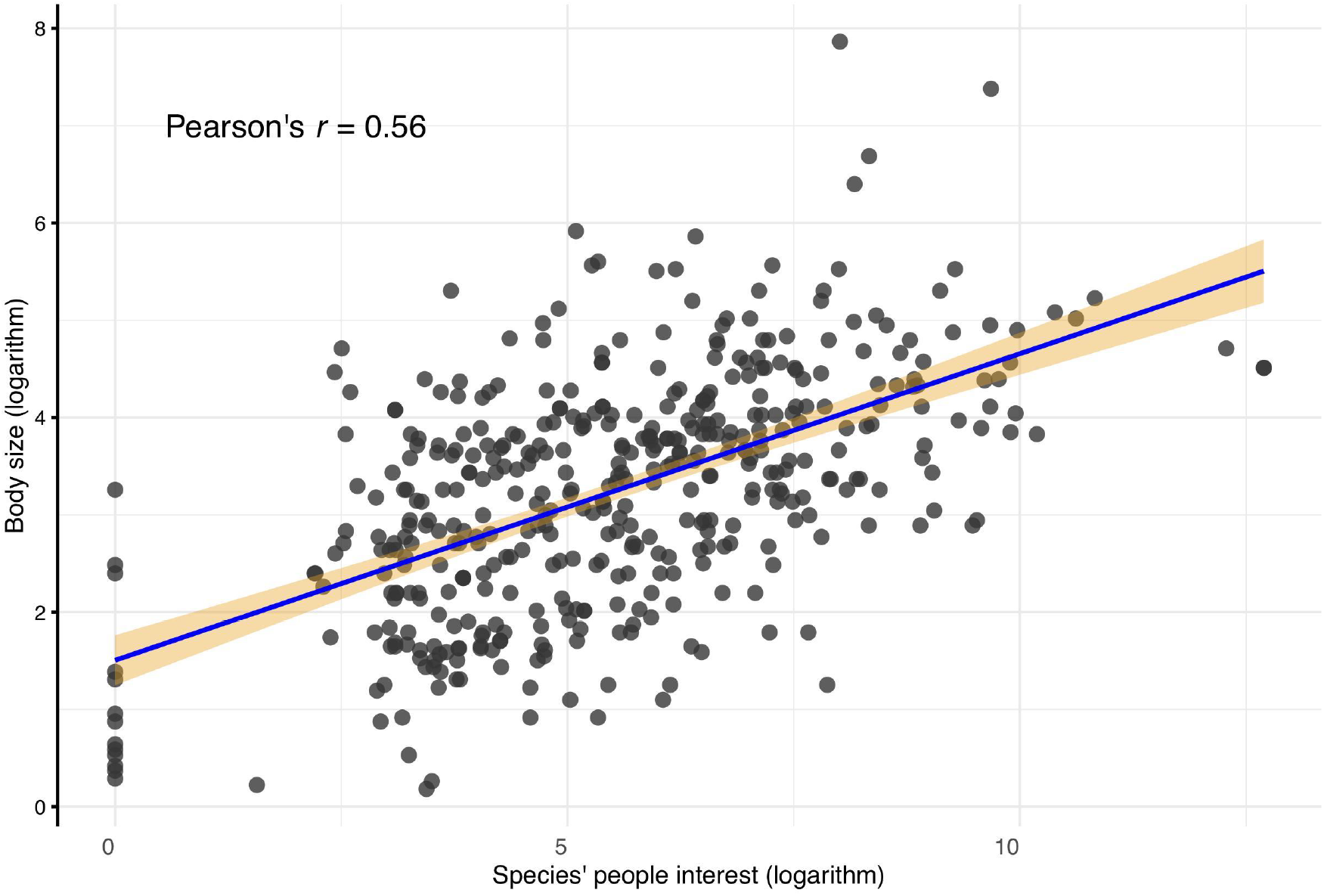
Relationship between body size and species’ online popularity.

## References

Cardoso, P. 2012. Habitats Directive species lists: Urgent need of revision. - Insect Conserv. Divers. 5: 169–174.

Cardoso, P. and Leather, S. R. 2019. Predicting a global insect apocalypse. - Insect Conserv. Divers. 12: 263–267.

Cardoso, P. et al. 2011. The seven impediments in invertebrate conservation and how to overcome them. - Biol. Conserv. 144: 2647–2655.

Cardoso, P. et al. 2020. Scientists’ warning to humanity on insect extinctions. - Biol. Conserv. 242: 108426.

Carmona, C. P. et al. 2020. Mapping extinction risk in the global functional spectra across the tree of life. - bioRxiv: 2020.06.29.179143.

Chapron, G. et al. 2014. Recovery of large carnivores in Europe’s modern human-dominated landscapes. - Science 346: 1517–1519.

Chichorro, F. et al. 2019. A review of the relation between species traits and extinction risk. - Biol. Conserv. 237: 220–229.

Chichorro, F. et al. 2020. Species traits predict extinction risk across the Tree of Life. - bioRxiv: 2020.07.01.183053.

Clark, J. A. and May, R. M. 2002. Taxonomic Bias in Conservation Research. - Science 297: 191–192.

Commission, E. 2020. Communication from the commission to the european parliament, the council, the european economic and social committee and the committee of the regions: EU Biodiversity Strategy for 2030 Bringing nature back into our lives.

Davies, T. et al. 2018. Popular interest in vertebrates does not reflect extinction risk and is associated with bias in conservation investment. - PLoS One 14: e0212101.

de Jong, Y. et al. 2014. Fauna Europaea - All European animal species on the web. - Biodivers. Data J. 2: e4034.

Dinerstein, E. et al. 2019. A Global Deal For Nature: Guiding principles, milestones, and targets. - Sci. Adv. 5: eaaw2869.

Dirzo, R. et al. 2014. Defaunation in the Anthropocene. - Science 345: 401–406.

Eisenhauer, N. et al. 2019. Recognizing the quiet extinction of invertebrates. - Nat. Commun. 10: 1–3.

Fukushima, C. et al. 2020. Global wildlife trade permeates the Tree of Life. - Biol. Conserv., 247: 108503.

Jarić, I. et al. 2020. iEcology: Harnessing Large Online Resources to Generate Ecological Insights. - Trends Ecol. Evol. 35: 630–639.

Ladle, R. J. et al. 2016. Conservation culturomics. - Front. Ecol. Environ. 14: 269–275.

Lewis, S. L. and Maslin, M. A. 2015. Defining the Anthropocene. - Nature 519: 171–180.

Mammides, C. 2019. European Union’s conservation efforts are taxonomically biased. - Biodivers. Conserv. 28: 1291–1296.

Moser, D. et al. 2016. Weak agreement between the species conservation status assessments of the European habitats directive and red lists. - Biol. Conserv. 198: 1–8.

Müller, A. et al. 2020. Evaluating and expanding the European Union’s protected-area network toward potential post-2020 coverage targets. - Conserv. Biol. 34: 654–665.

Rounsevell, M. D. A. et al. 2020. A biodiversity target based on species extinctions. - Science 368: 1193–1195.

Stork, N. E. 2018. How Many Species of Insects and Other Terrestrial Arthropods Are There on Earth? - Annu. Rev. Entomol. 63: 31–45.

## Supplementary literature

Aulagnier, S. et al. 2018. Mammals of Europe, North Africa and the Middle East.

Correia, R. A. 2019. Google trends data need validation: Comment on Durmuşoğlu (2017). - Hum. Ecol. Risk Assess. 25: 787–790.

Froese, R. and Pauly, D. 2000. FishBase 2000: concepts, design and data sources. - ICLARM. IUCN 2020. Red List version 2020–1.

Ladle, R. J. et al. 2016. Conservation culturomics. - Front. Ecol. Environ. 14: 269–275.

Lüdecke, D. et al. 2020. performance: Assessment of Regression Models Performance.: R package version 0.4.4.

Pinheiro J, Bates D, DebRoy S, S. 2019. Linear and Nonlinear Mixed Effects Models. - R Packag. version 3.1–140

R Core Team 2018. R: A Language and Environment for Statistical Computing.

Speybroeck, J. et al. 2016. Field guide to the amphibians and reptiles of Britain and Europe. - Bloomsbury publishing.

Svensson, L. et al. 2010. Collins bird guide 2nd edition. - British Birds.

Weiss, K. C. B. and Ray, C. A. 2019. Unifying functional trait approaches to understand the assemblage of ecological communities: synthesizing taxonomic divides. - Ecography 42: 2012–2020.

Zuur, A. F. and Ieno, E. N. 2016. A protocol for conducting and presenting results of regression-type analyses. - Methods Ecol. Evol. 7: 636–645.

Zuur, A. F. et al. 2009a. A protocol for data exploration to avoid common statistical problems. - Methods Ecol. Evol. 1: 3–14.

Zuur, A. F. et al. 2009b. Mixed Effects Models and Extensions in Ecology with R Extensions in Ecology with R Mixed Effects.

